# A quantitative systems pharmacology analysis of KRAS G12C covalent inhibitors

**DOI:** 10.1101/153635

**Authors:** Edward C. Stites, Andrey S. Shaw

## Abstract

The KRAS oncogene is the most common, activating, oncogenic mutation in human cancer. KRAS has proven difficult to target effectively. Two different strategies have recently been described for covalently targeting the most common activating KRAS mutant in lung cancer, KRAS G12C. Previously, we have developed a computational model of the processes that regulate Ras activation and this model has proven useful for understanding the complex behaviors of Ras signaling. Here, we use this model to perform a computational systems pharmacology analysis of KRAS G12C targeted covalent inhibitors. After updating our model to include Ras protein turnover, we verified the validity of our model for problems in this domain by comparing model behaviors with experimental behaviors. The model naturally reproduces previous experimental data, including several experimental observations that were interpreted as being contrary to conventional wisdom. Overall, this suggests that our model describes the Ras system well, including those areas where conventional wisdom struggles. We then used the model to investigate possible strategies to improve the ability of KRAS G12C inhibitors to inhibit Ras pathway signaling. We identify one, as of yet unexplored mechanism, that, if optimized, could improve the effectiveness of one class of KRAS inhibitor. We also simulated resistance to targeted therapies and found that resistance promoting mutations may reverse which class of KRAS G12C inhibitor inhibits the system better, suggesting that there may be value to pursuing both types of KRAS G12C inhibitors. Overall, this work demonstrates that systems biology approaches can provide insights that inform the drug development process.

## Introduction

Human cancer cells, such as those from pancreatic cancer, colorectal cancer, and lung cancer, commonly include somatically acquired *KRAS* mutations. KRAS is a GTPase that binds to guanine nucleotides GDP and GTP with high affinity. The GTP bound form of KRAS is considered the active form, and downstream signaling effectors specifically bind to the GTP bound form of KRAS. The cancer promoting KRAS mutations most commonly occur at codon 12, 13, or 61, and result in increased levels of GTP-bound KRAS which in turn promotes downstream signaling. It has long been believed that drugs with the ability to block aberrant KRAS signaling would benefit cancer patients.

A major advance in the targeting of KRAS has been made with the development of molecules that covalently and irreversibly bind to the cysteine residue of the KRAS G12C mutant [1-4]. The KRAS G12C mutant is particularly common in lung cancers. Selectivity of a drug for the cysteine in this mutant protein should result in specific targeting of KRAS G12C mutant containing cancer cells. Compounds that covalently interact with the codon 12 cysteine can be grouped into two different classes based upon which pocket the molecule settles into after bonding with cysteine. The compounds described by the Gray laboratory rest in the nucleotide pocket in place of a guanine nucleotide. G12C inhibitors that reside in the nucleotide pocket will be referred to as NP-inhibitors (or NPI). The compounds initially described by the Shokat laboratory rest in a distinct pocket, the “switch two pocket” that was previously unappreciated [1]. Compounds using this strategy will be referred to as SIIP-inhibitors (or SIIPI). The originally described NP-inhibitors and SIIP-inhibitors were not able to reliably target KRAS G12C in cell culture. A new SIIP-inhibitor, ARS-853, has been described that has much improved binding properties and displays KRAS G12C inhibition in cell culture [5].

Of note, the new SIIP-inhibitor has been effectively used to probe KRAS signal regulation. In studies published earlier this year, it was demonstrated that ARS-853 binds specifically to the GDP-bound form of KRAS [5, 6]. Oncogenic KRAS mutants are generally considered to be “constitutively active” and active cycling between GTP and GDP states has not been widely appreciated [4-8]. These studies also demonstrated a dependency upon the activity of RAS positive regulators known as GEFs (<u>G</u>uanine nucleotide <u>E</u>xchange <u>F</u>actor<u>s</u>), which had also not been widely appreciated [5-8].

Previously, we developed a computational model of the processes that together regulate Ras signaling [9, 10]. The model has made multiple predictions about Ras signaling that went against conventional wisdom but have now been experimentally confirmed, e.g. [9, 11-17]. Here, we applied the model to the problem of targeting KRAS G12C. To do this, the model was updated to include protein turnover, the specific biochemical properties of the G12C mutant, and the described mechanisms of interaction for the two classes of inhibitors. Simulations of the updated model yield patterns of Ras activation that match well with previous experimental observations. Application of the model to NP-inhibitors finds that GEF loading is a property that could be optimized to make NP-inhibitors more effective. The model also suggests that mutations that promote resistance to SIIP-inhibitors may increase sensitivity to NP-inhibitors. Overall, this work demonstrates the potential of mechanistically accurate models of oncogenic signal regulation to contribute to quantitative systems pharmacology.

## Model Development and Analysis

### Model extension to include protein turnover and the G12C mutant

The details of the Ras model have been previously published and described in detail [9, 10, 18-20]. To study the G12C mutant targeted by a covalent inhibitor, the model was updated in two key ways. First, protein synthesis and degradation were added to the model. When the model was originally developed, we assumed that these processes would be slow relative to the processes that regulate steady-state Ras nucleotide binding and could therefore be ignored when studying oncogenic mutant activation. Since covalent inhibitors target G12C irreversibly (and could potentially increase degradation), we added a degradative process to the model. A schematic with the updates to the original model is shown in Figure 1. Briefly, the model now allows for degradation of Ras in any form, and production of Ras results in nucleotide free Ras which will rapidly bind to available cytoplasmic nucleotides upon modeled production. Protein degradation was modeled as a first order process with a half-life of approximately 24 hours (k_deg_=8×10^-6^/s) [21, 22]. Production was modeled to occur at a constant rate. This rate was set so the steady-state level of total Ras remains at the level of our original estimate for total cellular Ras. Values for these parameters are provided in Table I. The model is specified as a set of coupled, nonlinear, ordinary differential equations. Simulations in MATLAB were used to find steady-state levels of RasGTP and RasGTP-Effector complex, both of which were considered as measures of Ras pathway signaling [9].

**Figure 1.**
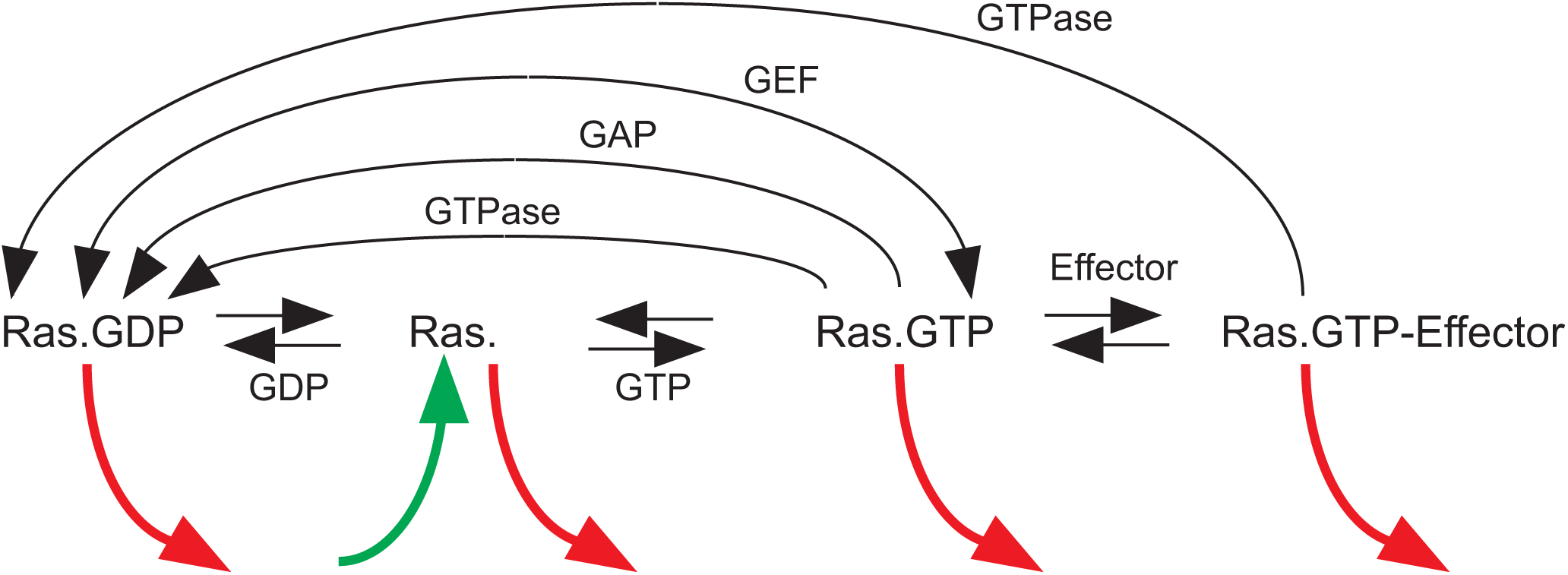
Modeled biochemical reactions of the Ras network including production and degradation. The original Ras model was based upon a dynamic equilibrium of Ras signaling states that followed from the biochemical reactions that influence Ras nucleotide binding state. This model was updated to include Ras protein production (green arrow) and degradation (red arrows). Parameters for these reactions are presented in Table I.

**Table I.**
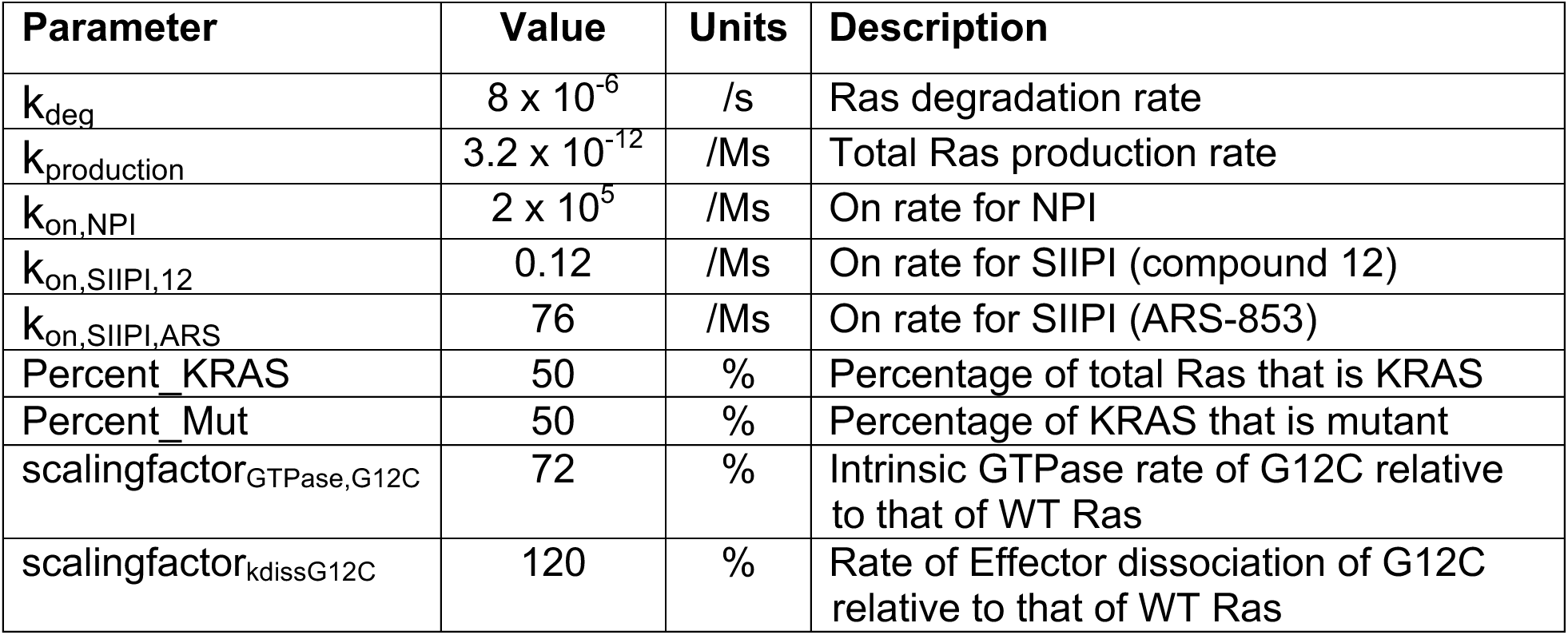
Parameters for the G12C model. Parameters for the expanded the Ras model that includes protein production and degradation, Nucleotide Pocket Inhibitors (NPI), Switch II Pocket Inhibitors (SIIPI), and the G12C mutant.

We also updated the model to include the G12C mutant. Of note, oncogenic mutants at codon 12, 13, and 61 are biochemically similar [23]. However, there are subtle differences between different oncogenic mutants that, in some cases, can have large effects [11, 24]. For that reason, we extended the model to include G12C by incorporating recently published data [25]. The parameters of the G12C mutant that vary from WT are the intrinsic GTPase rate and the effector binding constant [25]. The values used in our model are presented in Table I.

### Tests of the extended model with turnover and the G12C mutant

We first evaluated the behavior of the updated model by examining the predicted level of RasGTP and RasGTP-Effector complex for a cell with either G12D, G12V, G12C, or F28L mutations, as well as for a cell with all wild-type RAS. For these conditions, we modeled 25% of total RAS as mutant and 75% as wild-type, such as might occur if 50% of the RAS protein in the cell was KRAS, the other 50% NRAS and HRAS, and one of the KRAS alleles was mutated. Model predictions for levels of RasGTP and RasGTP-Effector were essentially the same whether or not Ras turnover was included (Figure 2). This confirmed the validity of our original assumption that turnover could be ignored for predicting behavior of oncogenic mutants in the absence of covalent inhibitors.

**Figure 2.**
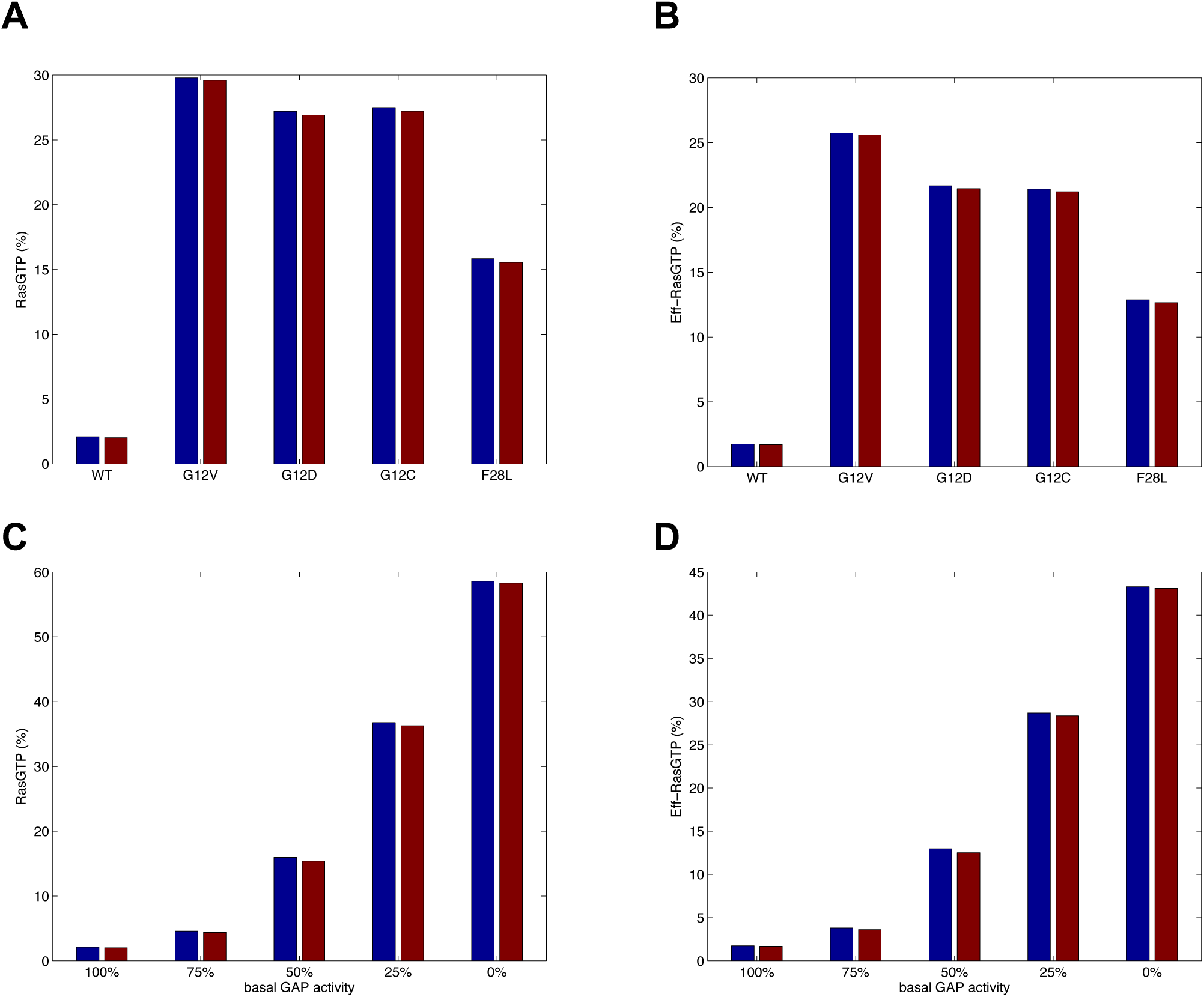
Predicted Ras signaling for oncogenes and tumor suppressors. The updated model diagrammed in Figure 1 was compared to the original model for predictions of Ras pathway activation due to oncogenic mutation and the tumor suppressor activity loss. Figures present computational predictions of the proportion of total Ras bound to GTP (A and C) and the proportion of total effector bound to RasGTP (B and D). A and B are for networks with all WT Ras, or with WT Ras and a mutant form of Ras (G12V, G1D, G12C, or F28L). C and D are for the Ras network with the indicated proportion of total basal GAP activity lost, such as might occur for a deletion or mutation to tumor suppressor NF1. Predictions with the original model are in blue, those with the updated model are in maroon.

We also investigated the behavior of the updated model by considering the loss of the tumor suppressor protein neurofibromin (NF1). Neurofibromin is a Ras <u>G</u>TPase <u>A</u>ctivating <u>P</u>rotein (Ras GAP). Ras GAPs act as negative regulators of RasGTP by promoting the conversion of RasGTP to RasGDP. Germline absence of a single copy of neurofibromin results in increased RasGTP, and loss of both copies of neurofibromin results in further increases in RasGTP [26]. NF1 is also one of the most commonly mutated driver genes in human cancer [27], where one or both copies can be mutated, deleted, and/or silenced.

Modeling 100% and 50% loss of total Ras-GAP in both the model with turnover and without turnover resulted in similar levels of RasGTP and RasGTP-Effector complex. We also considered fractions of total basal GAP loss other than 50% and 100% to consider conditions where other GAPs contribute to RasGTP homeostasis [28]. We considered 25% loss and 75% loss to span the range of levels of GAP activity, and found similar levels of RasGTP and RasGTP-Effector when the model with turnover was compared to the model without turnover. Overall, these simulations found that updating our model to include turnover results in essentially identical predictions for steady-state levels of Ras signal, whether caused by oncogenes or by loss of tumor suppressor genes.

### Extension of the model to covalent G12C inhibitors

We next extended the model to include the two types of KRAS G12C targeted covalent inhibitors (Figure 3). Our turnover model posits that NP-inhibitors bind to nucleotide-free G12C mutant irreversibly, consistent with the manner in which NP-inhibitors have been described to date [2, 3]. The SIIP-inhibitors are reported to bind to GDP bound G12C mutant (and presumably nucleotide free G12C mutant, as well), to bind irreversibly, and to prevent the inhibitor bound protein from exchanging GDP for GTP [5, 6]. We modeled SIIP-inhibitors with these same activities. Table I includes the reaction parameters for NP-inhibitors and SIIP-inhibitors. Of note, the reported difference between the original SIIP-inhibitor referred to as compound 12 [1] and the new SIIP inhibitor ARS-853 [5] are manifest exclusively in the on-rate of drug binding.

**Figure 3.**
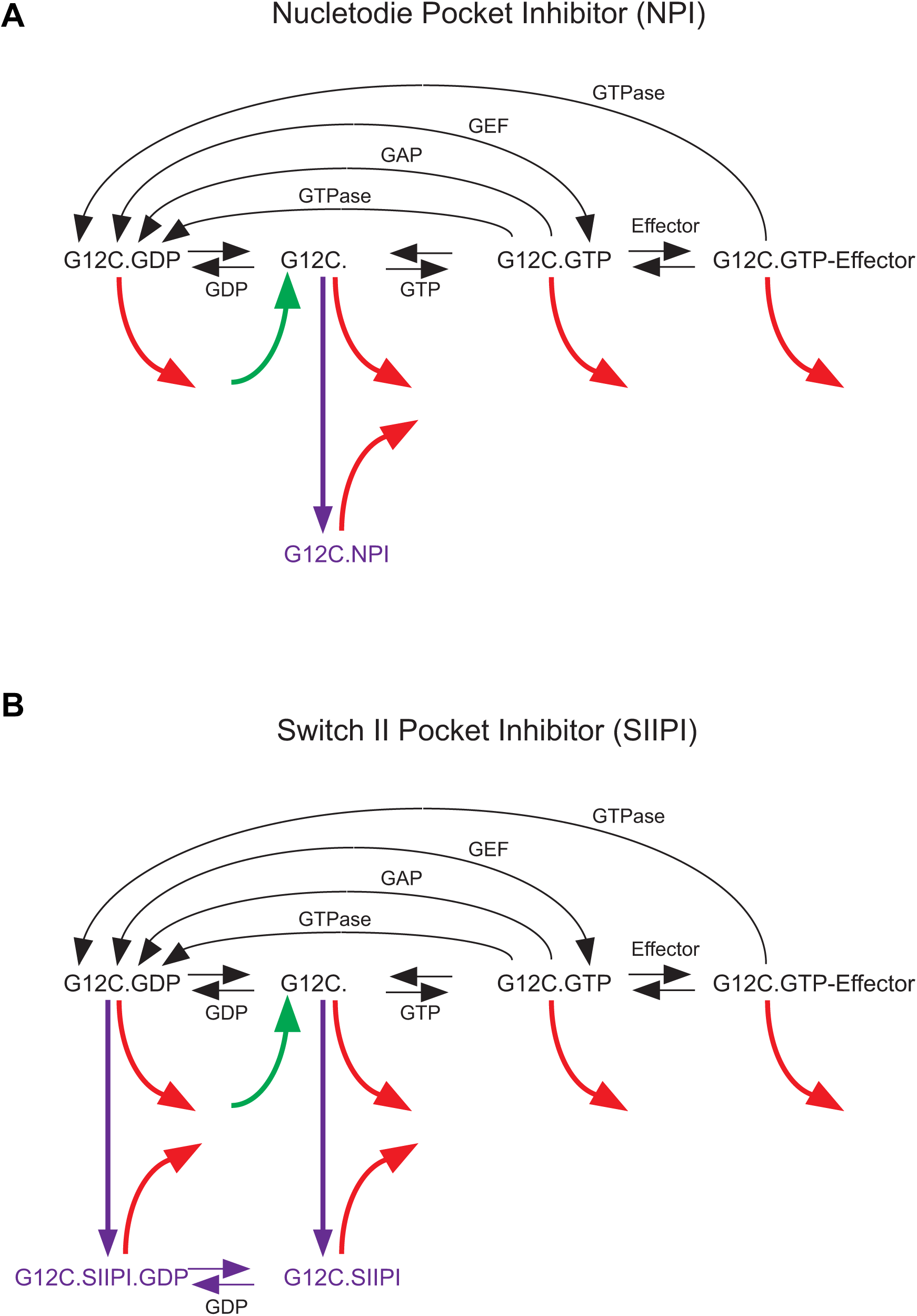
Modeled biochemical reactions of KRAS G12C targeted covalent inhibitors. The Ras model was further updated to include the two classes of G12C inhibitors. NPI bind to nucleotide free KRAS G12C, are subject to degradation, and are incapable of binding other nucleotides once drug is bound. SIIPI bind to GDP bound KRAS G12C and nucleotide free KRAS G12C, are subject to degradation, and GTP is not permitted to bind to drug bound G12C. All other interactions with GEFs, GAPs, and Effectors are assumed to be impermissible, consistent with available data.

### Model Validation: Testing the model’s ability to reproduce the behaviors of G12C inhibitors

We then simulated dose responses for NPI and SIIPI (Figure 4). NPI and SIIPI would appear similarly effective at steady-state if turnover was not modeled. However, when turnover was included in the model, the SIIP-inhibitor (ARS-853 compound) was clearly superior to the other SIIPI and to the NPI. In this more physiological case, the maximal amount of drug that will bind to KRAS G12C is limited by rate of each compound binding to KRAS G12C which in the case of the NPI is restricted by the infrequency of nucleotide free mutated KRAS. This highlights the importance of considering turnover when evaluating covalent inhibitors.

**Figure 4.**
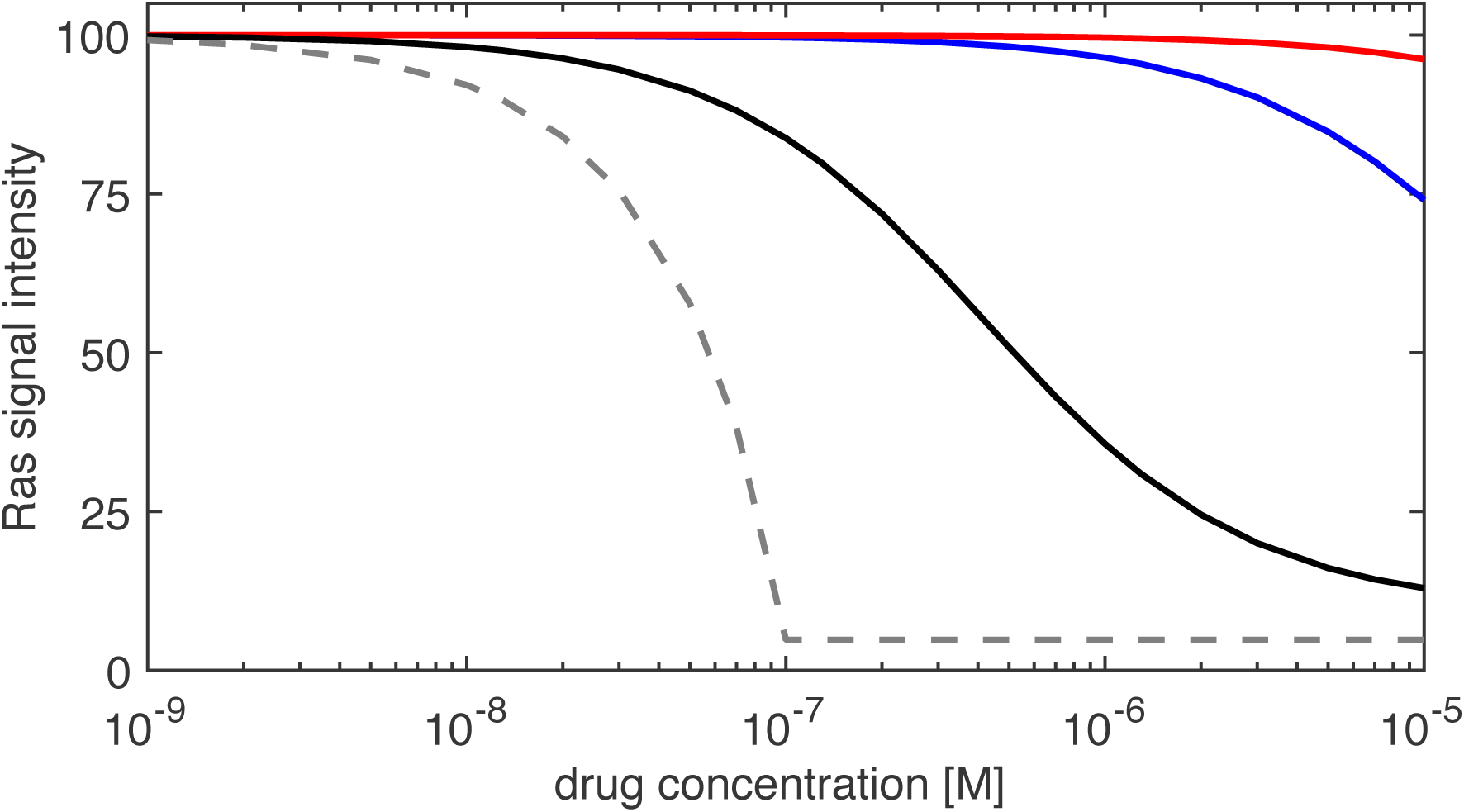
Simulated G12C inhibitor dose responses. The updated Ras model was used to simulate dose responses for NPI (blue) and SIIPI (red and black). For SIIPI, simulated dose responses were generated using both an on rate consistent with compound 12 described by Ostrem et al (red) and an on rate consistent with ARS-853 described by Patricelli et al (black). For comparison, the dose response for covalent inhibitors in the case where protein turnover is not modeled is included (dashed line).

Simulated behaviors of these drugs are consistent with their described abilities to inhibit KRAS G12C containing cancer cell lines, suggesting that our model includes the aspects of Ras biology needed to study KRAS G12C targeted covalent inhibitors. Experimentally, only the ARS-853 compound is consistently effective at inhibiting KRAS G12C lung cancer cells. Lito et al reported ARS-853 has an IC50 of approximately 2.5 microM [6]; Patricelli et al reported IC50 values ranging from 1 microM to 2 microM [5]. In our simulations, the range of greatest change in Ras signal was in the low μmolar range, with approximately 50% of Ras inhibition occurring at 1 μM, roughly mirroring the experimental observations. Lito et al. also reported an approximately 95% reduction of Ras signal when 10 μM of ARS-853 is applied, which is also comparable to the reductions in Ras signal found in our simulations. It is worth restating that these inhibitors are covalent and therefore do not have a K_d_; they only have a kinetic reaction rate constant for the binding reaction. This highlights that the agreement between the model and experiment is not due to assigning a K_d_ for the drug in this range (as no such K_d_ exists to be assigned), but rather suggests that the agreement reflects an accurate portrayal of the system.

### Model Validation: Testing the model’s ability to reproduce EGFR modulation of Ras inhibition

Two studies published earlier this year demonstrate that the NP-inhibitor ARS853 does not bind to GTP bound KRAS G12C, but rather binds to GDP bound KRAS [5, 6]. This suggests that oncogenic mutants exist in both GDP and GTP forms and this interpretation was considered inconsistent with conventional wisdom about Ras biology [5-8]. However, we note that such a dynamic equilibrium has long been implied by the available data on Ras nucleotide binding, and such a dynamic equilibrium was a part of our original model developed a decade ago [9]. Our previously published model predicted that approximately 80% of the mutant is bound to GTP and approximately 20% of the mutant is bound to GDP [9], consistent with data presented in 2016 by Lito et al. [6]. Thus, we next considered whether our model can reproduce other biological experiments used to explore the interplay between SIIPI and the dynamic equilibrium of nucleotide binding.

Multiple experiments probing the effects of EGFR activation (or inhibition) on the inhibition of RAS signaling by SIIPI provide additional test situations to validate our model. The activation of EGFR delays the kinetics of inhibition, and the inhibition of EGFR expedites the kinetics of inhibition [6]. This suggests that the dynamics of RAS activation can modify the effect of the drugs. To simulate these experiments, we modeled EGFR inhibition by decreasing the concentration of basally active GEFs in our model and we modeled EGFR activation by increasing the concentration of basally active GEFs. We found that our model reproduces the kinetic changes that follow from EGFR activation and inhibition (Figure 5A). Of note, the changes in total RasGTP that our model predicts were also observable in the previous experiments. We also considered how EGFR/RTK activation would influence the ARS-853 dose response. We found that the SIIP inhibitor dose response would shift to require higher levels of drug in situations of EGFR activation and to require less drug for situations of EGFR inactivation (**Figure 5C,D**), consistent with the measured dose responses [5, 6].

**Figure 5.**
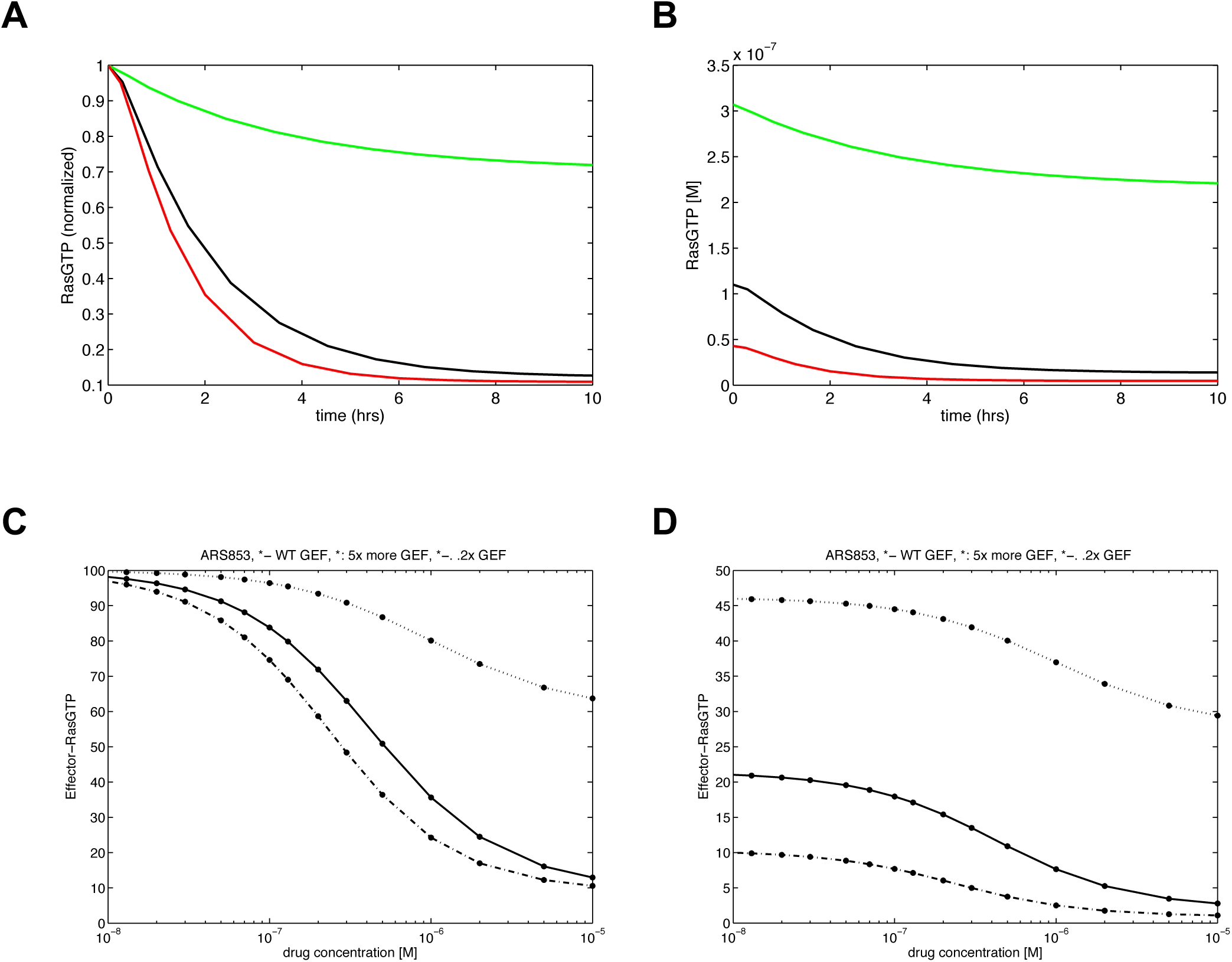
The Ras model predicts how RTK activation and inhibition influences ARS-853 inhibition. Simulations of Ras G12C network inhibition by the SIIPI ARS-853 were performed, including conditions of upstream activation and upstream inhibition. Simulations of the kinetics of the inhibition found increased GEF activity (upstream activation) delays the kinetics of inhibition (green, A and B) while decreased GEF activity results in faster inhibition (red, A and B). Kinetics for the basal model (no change in GEF activity) is shown in black. (A) is normalized to the total amount of Ras signal in each case, while (B) shows total RasGTP in each case. The steady-state dose responses of ARS-853 in these same conditions of increased or decreased GEF activity were also found through model simulations (C and D). Increased GEF activity (dotted line) decreased GEF activity (dashed line), basal GEF activity (solid line). (C) presents the data scaled to maximal amount of Ras signal, while (D) shows the total amount of Ras signal.

### Model Validation: Testing the model’s ability to simulate resistance promoting mutations

In their characterization of the behaviors of the SIIPI, Lito et al engineered specific KRAS mutations that had both the G12C mutation as well as another mutation at a separate KRAS residue to introduce additional biochemical defects [6]. They considered secondary mutations that would be anticipated to potentiate nucleotide exchange (e.g. Y40A, N116H, or A146V) and that would be anticipated to impair GTPase activity (e.g. A59G, Q61L, or Y64A). Lito et al hypothesized and demonstrated that these secondary mutations would have reduced sensitivity to ARS-853.

We evaluated our model’s ability to reproduce these experiments. To simulate combinations of mutations within the same Ras protein, such as combinations of G12C with secondary mutations that are anticipated to reduce the intrinsic GTPase rate (e.g. KRAS G12C/A59G) or secondary mutations that are anticipated to increase the spontaneous exchange of nucleotide (e.g. KRAS G12C/A146V) we maintained the G12C parameters and adjusted only the rate constant(s) that were anticipated to change as a consequence of the secondary mutation. Our model simulations showed that increasing impairment of GTPase activity should make the mutant increasingly less sensitive to ARS853 (Figure 6A) and that faster cycling secondary mutants should also be less sensitive to ARS853 (Figure 6B); both model behaviors are consistent with the previous experimental observations [6].

**Figure 6.**
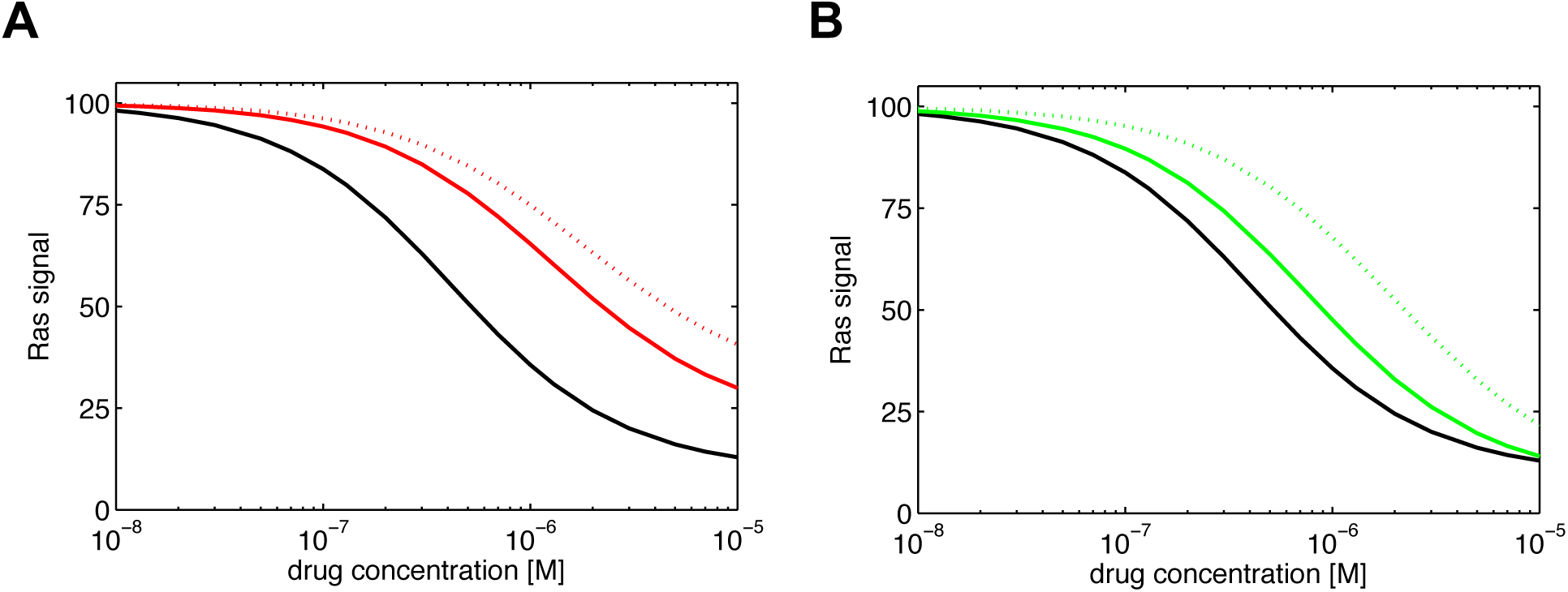
Modeled secondary mutations to G12C and ARS-853 dose response. Simulations were performed to evaluate the conditions where Lito et al engineered secondary mutations into KRAS G12C that impaired GTPase activity and that altered the rate of nucleotide exchange. (A) Decreased rates of intrinsic GTPase activity are predicted to result in a decreased response to ARS-853. Solid red: 10x slower GTPase activity than G12C, dotted red: 100x slower GTPase activity than G12C. (B) Increased intrinsic nucleotide dissociation is predicted to result in a decreased response to ARS-853. Solid green: 10x faster, dashed green: 100x faster, both relative to G12C.

### Model validation: Conclusion

We considered several different experiments that tested the inhibition of Ras signaling with the newly developed G12C inhibitors. We found that the model could readily reproduce experimentally observed behaviors, suggesting that our model is valid for problems involving G12C inhibitors. Additionally, it is worth noting that multiple experimental behaviors that were interpreted as contrary to conventional wisdom could have been predicted *a priori* had our readily available model first been applied to these problems.

### Analysis of NP Inhibitors

We next used the model to investigate NP-inhibitors. We considered why NPI are less effective than SIIPI (Figure 7). A key step for the drug, as described, is binding to a non-occupied nucleotide binding pocket [3]. A non-occupied nucleotide binding pocket is likely to be rarely encountered within the cell due both to the high affinity of the pocket for guanine nucleotides and also the high concentration of guanine nucleotides within the cell.

**Figure 7.**
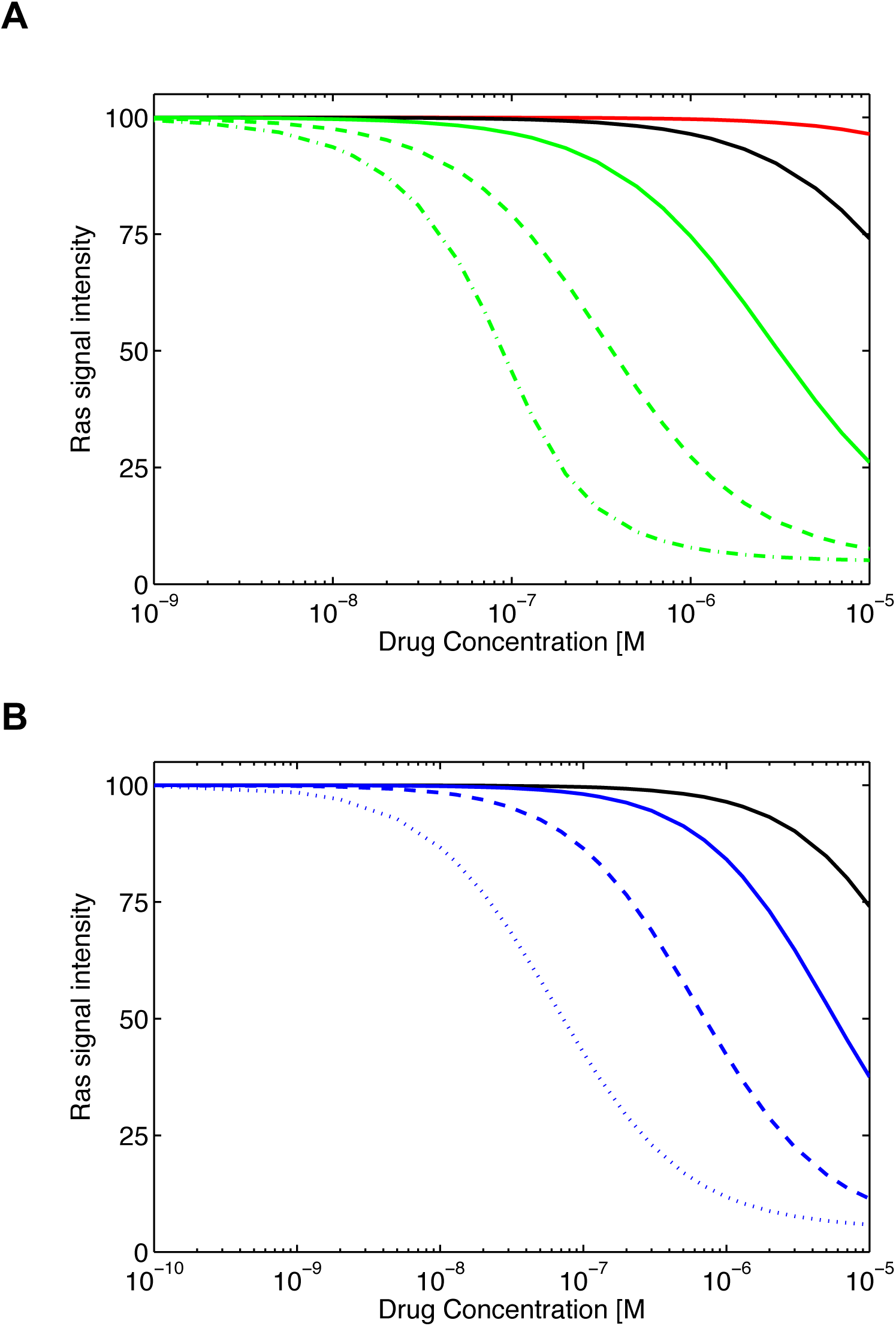
Simulations identify methods to improve NPI effectiveness. The NPI inhibitor described by Lim et al and Hunter et al was studied to determine how optimizing different biochemical rate constants might augment the effectiveness of NPIs. (A) An increased and decreased on rate of the compound was modeled: red: 10x slower, solid green: 10x faster, dashed green: 100x and 1000x faster. (B) The ability of GEFs to load the NPI, comparable to GEF loading of nucleotides, was considered. Solid blue: GEF loading of NPI proportional to the abundance of NPI and nucleotides. Dashed blue: GEF loading of NPI favoring NPI over nucleotides by a factor of ten over their proportional abundance. Dotted blue: GEF loading of NPI favoring NPI over nucleotides by a factor of one hundred over their proportional abundance.

We initially modeled NPI to bind the Ras nucleotide pocket with an on rate similar to that for nucleotides (k_on_ = 2×10^5^/Ms). We next considered on rates that were one, two, and three orders of magnitude faster to evaluate how much better inhibition would be if the on rate could be increased. Computational simulations suggested that a significant enhancement of the on rate would result in a much-improved dose response, even when considering that nucleotide free Ras protein is very limited within the cell (Figure 7A). However these levels are approaching the theoretical diffusion limit [29], suggesting that it would not be possible to engineer a better NPI by only optimizing the forward reaction rate constant.

We considered whether GEFs could facilitate the loading of an NPI. The authors of the NP-inhibitor studies did not address whether or not Ras GEFs could facilitate the loading of their compound. Their *in vitro* studies with recombinant protein did not include Ras GEFs, and much of the conventional wisdom on oncogenic mutants is that GEFs do not contribute to oncogenic signaling [8]. However, our previous modeling studies of Ras suggest that a small level of basal GEF activity is needed to explain the experimental data of basal nucleotide exchange, and that GEF inhibition should result in less oncogenic Ras signaling [9]. We therefore considered the potential effects of GEF loading on NP-inhibitors.

Our simulations suggest that GEF loading could be an important variable for NPI. If GEF loads the G12C inhibitor as well as it loads nucleotide (e.g. it loads the G12C inhibitor like GTP and GDP, and at a rate that is proportional to the abundances of drug and nucleotides), our simulations suggest that there would be a modest increase in the ability of NP-inhibitors to target the G12C mutant (Figure 7B, blue solid line). If GEF loading could favor drug loading by a factor of ten to one hundred over nucleotide, the effectiveness of this class of drug is predicted to be much higher (Figure 7B, blue dotted and dashed lines). This computational analysis therefore suggests that efforts to optimize NP-inhibitors should evaluate candidates for whether or not GEFs facilitate their loading. Of note, the predicted levels of optimized (GEF-facilitated) NPI (oNPI) needed to inhibit 50% of the Ras signal were on the order of 1 μM, much smaller than the near mM levels of nucleotide that are found within the cell.

### Analysis of optimized NPI on resistance promoting mutations

We next considered how optimized NPI would respond to the mechanisms described to promote resistance to SIIPI. Our simulations found that increased intrinsic nucleotide cycling, such as what occurs with Y40A, N116H, and A146V compound mutations with G12C, could actually result in an increased sensitivity to NPI (Figure 8A). This suggests that there may be value in the continued development of optimized NPIs. In contrast, our simulations found that secondary mutations to KRAS that impair GTPase activity (Figure 8B) or mutations that increase RTK activation (Figure 8C) would both result in less sensitivity to NPI. This suggests that an optimized NPI would not be able to combat all potential forms of G12C acquired resistance. It is worth noting that although the optimized NPI was less effective than the SIIPI for a G12C mutant, the optimized NPI was more effective than the SIIPI for all three resistance promoting mutations.

**Figure 8.**
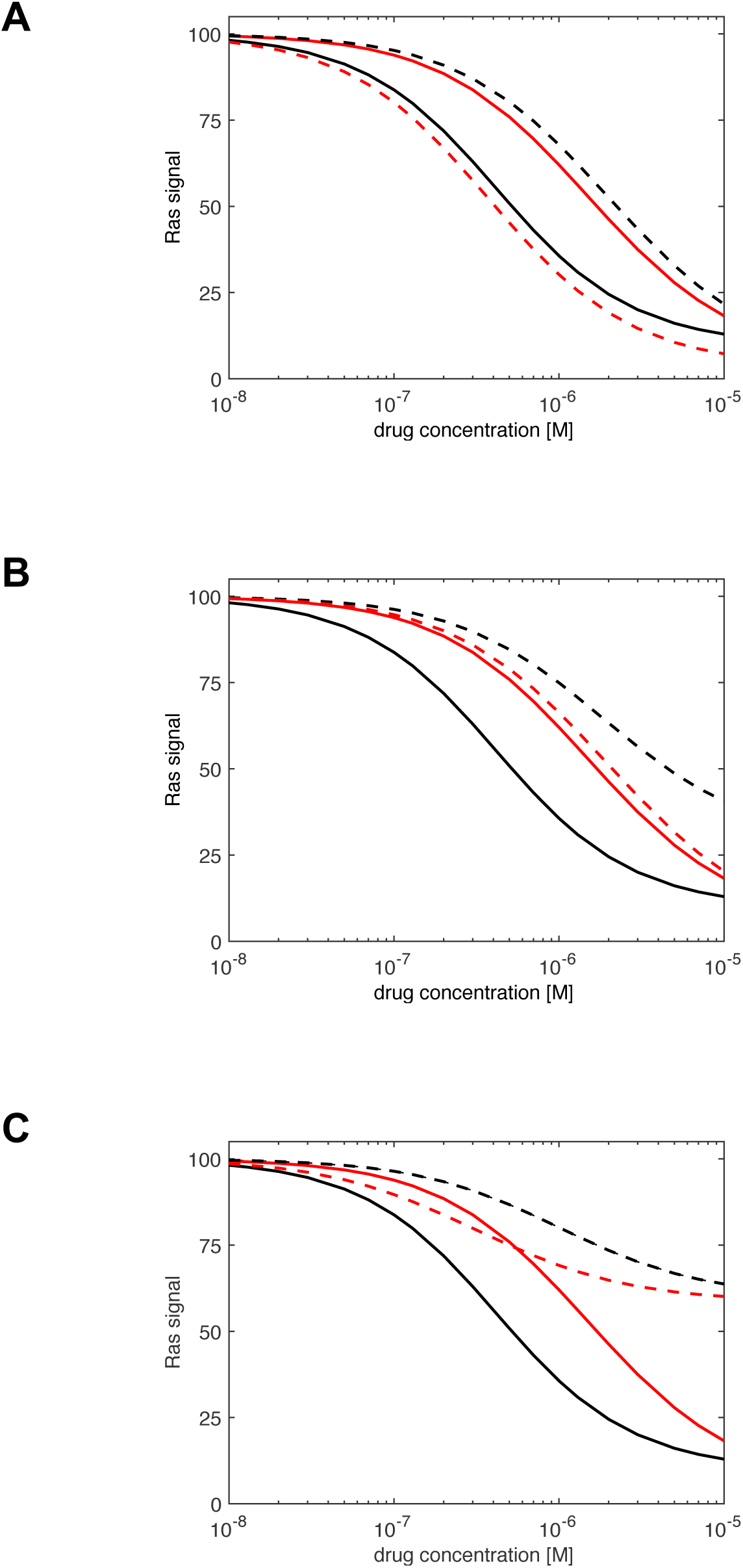
Simulations find SIIPI resistant promoting mutations can make NPIs the more effective KRAS G12C inhibitor. Potential SIIPI resistance promoting mutations were studied for both an optimized NPI and a SIIPI. The optimized NPI (oNPI) was assumed to have its loading facilitated by GEFs by a factor of four over the proportional abundance of NPI to total guanine nucleotides. (A) For the G12C mutant, the SIIPI ARS-853 (black solid) was predicted to be superior to the oNPI (red solid), but once a secondary mutation that caused increased nucleotide exchange was modeled, (100x over G12C) simulations found a decreased effectiveness of the SIIPI (black dashed), but a higher effectiveness for the oNPI (red dashed). (B) When a secondary mutation that causes decreased intrinsic GTPase activity was modeled (100x relative to G12C), both the SIIPI (black dashed) and oNPI (red dashed) were less effective. In the conditions of the secondary mutation, the oNPI became relatively more effective than the SIIPI. (C) When a secondary mutation increased RAS activation through increased GEF activation (e.g. as might happen with an RTK mutation), the oNPI (red dashed line) became more effective than the SIIPI (black dashed), although both classes of inhibitor are predicted to be overall less effective against mutations that result in increased GEF activation.

## Discussion

One interesting finding from the model is that it suggests NP-inhibitors could be more effective if GEFs can promote their loading into the nucleotide pocket. This possibility seems consistent with known GEF biology and needs to be considered experimentally. However, the ability of Ras GEFs to load NP-inhibitors has not, to the best our knowledge, been addressed to date. The reason it has not been addressed previously may be because it is widely believed that the increased activation of oncogenic Ras is independent of GEFs. Our work suggests that efforts to screen derivatives of the known NP-inhibitors and new NP-inhibitor candidates should include the evaluation of how well Ras GEFs can load the compound. This could be done experimentally by adapting the recombinant protein based assay used by Hunter et al [3] to also include a recombinant GEF and determining how NPI binding to RAS is enhanced by GEFs. Our work further suggests that compounds that are loaded by GEFs preferentially over nucleotides would be most beneficial.

Another interesting finding was that theoretically, the better inhibitor could change as secondary resistance mutations are acquired. For example, the modeled GEF-facilitated NP-inhibitor is inferior to the SIIP inhibitor with respect to inhibiting KRAS G12C signaling. However, the model suggests that the GEF-facilitated NP-inhibitor should be superior to the SIIP-inhibitor if a G12C mutant picks up a second mutation to the same allele that results in faster nucleotide dissociation.

A major challenge in developing targeted therapies for cancer is that it can be difficult to anticipate the response of a biological network to an inhibitor. Indeed, multiple experiments using KRAS G12C inhibitors have been interpreted as unexpected and contradictory to conventional wisdom. Computational systems biology models can provide an alternative viewpoint to expert opinion. It is possible to use a computational model to find the systems level behaviors that naturally emerge from the constituent reactions when the model is based upon the fundamental reactions of the network. We have done that here, and we demonstrate that multiple experimental observations that were interpreted as unexpected were foreseeable by computational modeling. We anticipate that models like this will be increasingly utilized to inform efforts to develop and use targeted therapies.

Altogether, we have applied a computational systems biology approach to the analysis of targeted covalent inhibitors. Specifically, we used a model of the biochemical reaction network that regulates Ras signals to study how two specific KRAS G12C oncogenic mutant inhibition strategies alter Ras signaling. Our model finds the behaviors that logically follow from what is known about Ras signaling and about these inhibitors. We find that our model naturally reproduces the experimentally observed behaviors of both NPI and SIIPI, including results that were widely perceived as unexpected and inconsistent with known Ras biology. Our model also suggests strategies to improve the effectiveness of one of these classes of inhibitors.

